# Discovery of Isobavachin, a natural flavonoid, as an Apolipoprotein E4 (ApoE4) structure corrector for Alzheimer’s disease

**DOI:** 10.1101/2025.01.03.631209

**Authors:** Sachin P. Patil, Bella Kuehn, Christina McCullough, Dean Bates, Hadil Hazim, Mamadou Diallo, Naomie Francois

## Abstract

Alzheimer’s disease (AD) is a progressive neurodegenerative disease characterized by extensive neurodegeneration and consequent severe memory loss. Apolipoprotein E4 (ApoE4) is the strongest genetic risk factor for AD, with its pathological effects linked to structural instability and altered interactions with lipids and other important disease proteins including amyloid beta (Aβ) and tau (τ). Therefore, correcting and stabilizing the ApoE4 structure has emerged as a promising therapeutic strategy for mitigating its detrimental effects. In this study, we investigated naturally occurring bioavailable flavonoids as ApoE4 stabilizers, focusing on their potential to modulate ApoE4 structure and function. Comprehensive investigation of a focused database using our integrated computational and experimental screening protocol led to the identification of Isobavachin as a potential corrector and stabilizer of ApoE4 structure. In addition, a few other bioavailable flavonoids with similar stabilizing properties were identified, albeit to a much lesser extent as compared to Isobavachin. The findings support the therapeutic potential of flavonoids as ApoE4 modulators and highlight Isobavachin as a lead candidate for further preclinical evaluation. These results provide new insights into the pharmacological targeting of ApoE4 and open avenues for the development of flavonoid-based, ApoE-directed therapies for AD.

## 1. Introduction

Alzheimer’s disease (AD) is a progressive neurodegenerative disorder and the most common form of dementia, posing an immense socioeconomic challenge with over 55 million people living with AD and other dementia worldwide [1]. Clinically, AD is characterized by significant memory loss, cognitive decline, and eventual loss of functional independence [2]. Pathologically, it involves the accumulation of extracellular amyloid-beta (Aβ) plaques and intracellular neurofibrillary tangles composed of hyperphosphorylated tau (τ) protein [3–6].

Numerous studies imply precedence of Aβ accumulation before other AD-related phenotypes including neurofibrillary τ tangle formation, reactive gliosis, activated microglia and complement pathways, severe inflammation, early synapse loss, and neurotransmitter deficit, leading in turn to irreversible cognitive impairment [7–9]. Accordingly, the “Amyloid Cascade Hypothesis” has been considered central to the AD pathogenesis, receiving major preclinical and clinical research attention so far [10]. In this context, two anti-Aβ monoclonal antibodies, viz. Aducanumab and Lecanemab, have recently received accelerated approvals by the United States Food and Drug Administration (FDA) [11,12]. Despite their effectiveness in reducing the brain Aβ accumulation, the therapeutic efficacy of these approved and other investigational Aβ-directed approaches on cognitive decline is unfortunately somewhat limited [13]. Moreover, the serious adverse effects including brain edema together with high costs plague the widespread use of these biological drugs against AD. As such, the European Medical Agency (EMA) declined to license Aducanumab across Europe citing lack of safety and clinical efficacy, and the manufacturer Biogen abandoned its commercial development completely in January 2024. Thus, despite decades of research, the therapeutic landscape for AD remains limited. Indeed, the current pharmacological approaches for AD include three cholinesterase inhibitors (Donepezil, Galantamine and Rivastigmine) and a N-methyl-D-aspartate (NMDA) receptor antagonist (Memantine), all of which have beneficial but only short-lived (lasting ∼ 6 months) effects in mediating only the symptoms of AD [13]. Therefore, there is a significant unmet need for novel, effective therapeutic approaches targeting alternative, fundamentally important disease pathways in AD.

Among such pathways, the role of Apolipoprotein E (ApoE), particularly its isoform-specific effect, is recently receiving significant traction as a promising avenue for AD intervention. ApoE, a 299-amino acid glycoprotein, is the primary lipid transporter in the brain, which exists in three major isoforms, viz. ApoE2, ApoE3, and ApoE4, differing by single amino acid substitutions at positions 112 and 158 [14]. These minor structural differences differentially impact ApoE risk for AD, with ApoE4 identified as the strongest genetic risk factor for late-onset AD, conferring a 10–15-fold increased risk in homozygous carriers compared to neutral ApoE3 [14]. The structural instability of ApoE4 profoundly impacts its biochemical properties including lipidation, receptor binding, protein stability, and aggregation propensity, exacerbating various neurodegenerative processes in AD [15–17]. Specifically, poorly lapidated ApoE4 impairs cellular lipid homeostasis, leading to exacerbation of Aβ deposition [18,19]. Also, ApoE4 binds less efficiently to its receptors including low-density lipoprotein receptors (LDLRs) and heparan sulfate proteoglycans (HSPGs), impairing lipid and Aβ clearance [20]. Furthermore, ApoE4 is less stable than ApoE3, leading to an increased tendency for self-aggregation, further contributing to its neuronal toxicity. Thus, stabilizing ApoE4 structure to prevent its pathological conformational changes has emerged as a promising therapeutic strategy against AD [21].

One of the well-studied approaches to stabilize ApoE4 structure is through targeting its “domain interaction”—a salt bridge interaction between the amino-terminal LDLR binding domain and the carboxy-terminal lipid binding domain [22]. This is directly attributed to the presence of Arginine at amino acid position 112 in ApoE4, instead of Cysteine112 observed in ApoE3, leading to the salt bridge interaction between Arginine61 in amino-terminal domain and Glutamate255 in carboxyl-terminal domain in ApoE4. This domain interaction in ApoE4 is found to lead to its closed structure as compared to more open structures of both ApoE2 and ApoE3 [23]. These ApoE structural differences, owing to the presence or absence of domain interaction, are responsible for their functional and related stability difference (ApoE2>ApoE3>ApoE4). Several small-molecule structure correctors that prohibit this domain interaction and convert ApoE4 to an ApoE3-like, more stable structure have been reported to mitigate the neurotoxic effects of ApoE4 [23,24].

More recently, an alternative approach in terms of stabilizing ApoE4 structure is reported by AbbVie Pharmaceuticals research team, involving small molecules that bind to an allosteric region of ApoE4 away from the domain interaction site [25]. Specifically, the Cysteine to Arginine substitution at 112 position was observed to induce long-distance conformational changes in ApoE4 protein. One such prominent change involves Tryptophan34 amino acid side chain assuming a “flip-out” orientation parallel to the ApoE4 axis as opposed to a “flip-in” orientation perpendicular to the axis of both ApoE2 and ApoE3 proteins (**Figure 1**). The small molecules identified by AbbVie team bound to a small but well-defined cavity formed by the rotation of the Tryptophan34 sidechain in ApoE4, stabilizing its structure and consequently suppressing inflammation in the neuronal cell culture [25]. Importantly, as seen in the crystal structure of ApoE4 in complex with the stabilizer molecule (**Figure 1d**), the ligand binding in the ApoE4 pocket is shown to shift the orientation of Tryptophan34 sidechain to that observed in both the free ApoE2 and ApoE3 structures (**Figures 1a-b**). Taken together, small-molecule correctors that can interact with Tryptophan34 residue in ApoE4 and induce its proper sidechain orientation may help in modifying the aberrant conformation of ApoE4 and in turn abolish its neurotoxic and other AD-relevant downstream effects.

**Figure 1.**
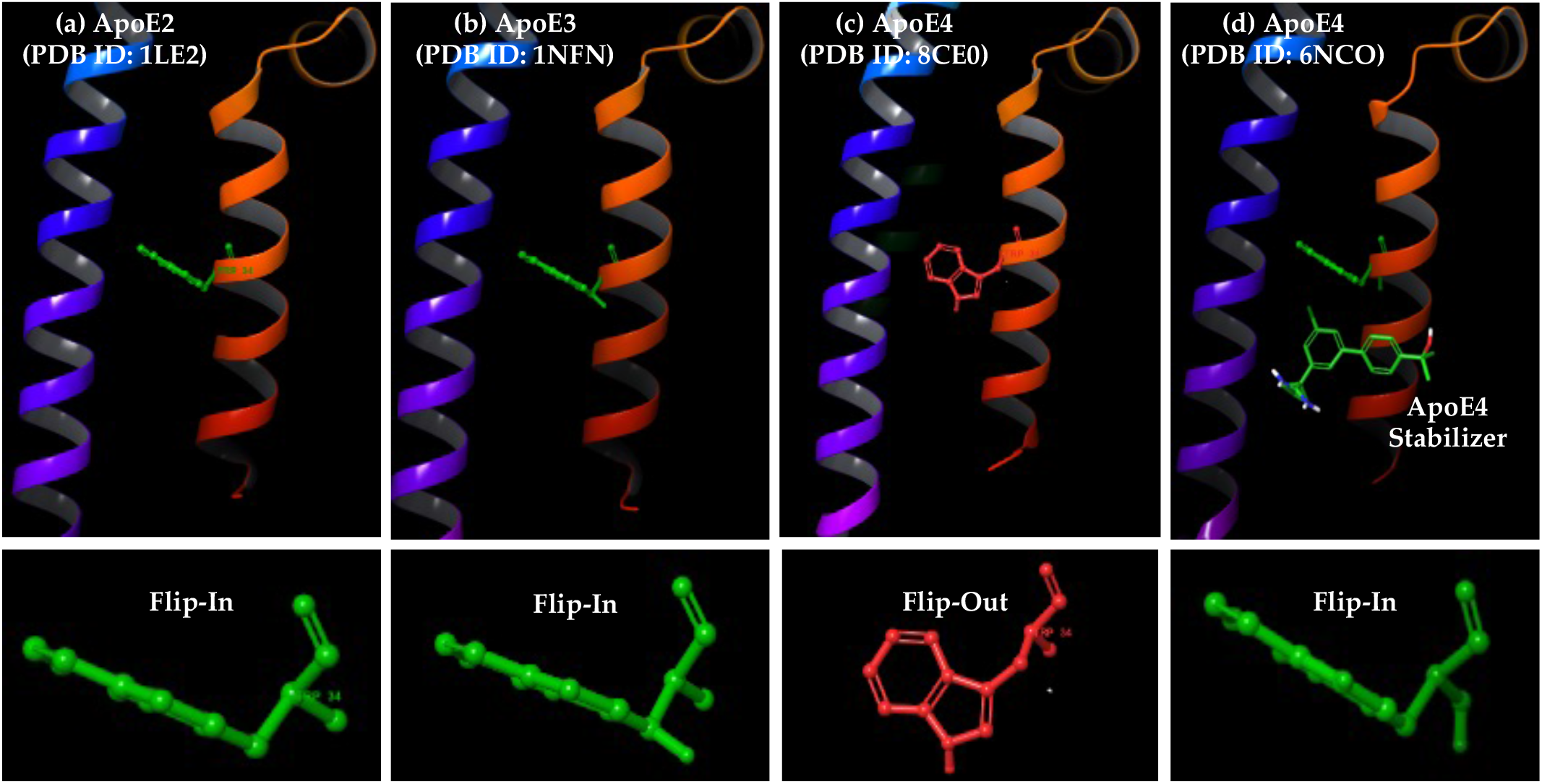
Crystal structures of different ApoE isoforms and orientation of Tryptophan34 residue. (a-c) Ribbon representation of ApoE2/E3/E4 apo structures, with Tryptophan34 shown as balls-and-sticks model; (d) The ApoE4 protein in complex with a small-molecule stabilizer from AbbVie Pharmaceuticals.

Therefore, given the critical role of ApoE4 in AD pathology, our present study focused on identifying small molecules that can stabilize the ApoE4 structure, thus reducing its aggregation propensity and restoring its function to resemble ApoE2/E3. Here, we utilized an integrated computational and experimental screening approach to systematically screen a focused database of bioactive small molecules for potential ApoE4 stabilizers. Specifically, the ensemble rigid-receptor docking approach was first used against two each apoas well as holo-ApoE4 crystal structures. This structure-based approach combined with our innovative consensus scoring analysis significantly enhanced the ranking for the known ApoE4 stabilizers and further led to the identification of several novel compounds with favorable molecular characteristics. These molecules were then subjected to flexible receptor docking and molecular dynamics studies, revealing Isobavachin, a bioavailable flavonoid, as a top virtual hit with the potential to bind to ApoE4 pocket and change the trytophan34 sidechain orientation towards resembling ApoE2/E3. The biophysical binding studies using Surface Plasmon Resonance (SPR) confirmed potent Isobavachin-ApoE4 binding with nanomolar range potency. Furthermore, we highlight a few additional flavonoids with similar but relatively weaker ApoE4 stabilizing potential.

In summary, our present work provides critical insights into the pharmacological targeting of ApoE4 and lays the groundwork for developing flavonoid-based therapies for AD. Isobavachin and other active flavonoids may serve as guiding points for the discovery and optimization of more potent, selective small-molecule stabilizers for ApoE4, one of the most promising therapeutic targets against AD. Importantly, the present results also validate potential viability of the combined computational and experimental screening protocol presented here for screening other available databases for identification of potentially novel ApoE4 stabilizers. Such molecules, by targeting ApoE4’s structural instability, may hold promise as disease-modifying therapies for AD.

## 2. Results

The present study involved use of combined structure- and ligand-based screening of a database containing drug-like compounds derived from the **T**raditional **C**hinese **M**edicine (**TCM**), viz. **D**atabase of **C**onstituents **A**bsorbed into the **B**lood and **M**etabolites of **TCM** (**DCABM-TCM**) [26]. The natural compounds are a precious source of novel chemical scaffolds and have been shown to be basis of 3/4^th^ of approved drugs worldwide during the past half century [27]. As such the DCABM-TCM database may prove to be a great resource of active natural products for AD drug development [28,29], offering potential ApoE4 stabilizers that are both bioactive and bioavailable. Therefore, virtual screening hits from the DCABM-TCM database were subjected to further computational analysis including the induced-fit docking and molecular dynamics simulations, followed by the experimental ApoE4 binding studies using SPR.

### 2.1. Selection of the ApoE structures for virtual screening

All three isoforms of ApoE are widely studied, with numerous 3D structures published using various methods, such as X-ray crystallography, **N**uclear **M**agnetic **R**esonance (**NMR**) and **E**lectron **M**icroscopy (**EM**). Specifically, UniProt database lists 22 X-ray structures, 5 NMR structures and 1 EM structure for various ApoE isoforms (UniProt ID: P02649). For the present study, we focused on the ApoE structures with high resolution of ≤ 2Å, leading to total 17 structures. Furthermore, we shortlisted only the natural, non-mutant structures, resulting in the final selection of three ApoE3 structures (PDB IDs: 1NFN, 1BZ4 and 1OR3), three ApoE4 structures (PDB IDs: 1B68, 8CE0 and 8CDY) and two holo-ApoE4 structures bound with small-molecule stabilizers (PDB IDs: 6NCN and 6NCO). Careful inspection of these eight ApoE crystal structures showed that one ApoE3 structure (PDB ID: 1OR3) showed Tryptophan34 side chain assuming a “flip-out” orientation that resembled ApoE4 structure, while one ApoE4 structure (PDB ID: 1B68) with Tryptophan34 side chain assuming a “flip-in” orientation that resembled ApoE3 structure (**Figure 2**). Therefore, we removed these two structures leaving two each of the free ApoE3, ApoE4 and holo-ApoE4 structures, for further computational analysis as described below.

**Figure 2.**
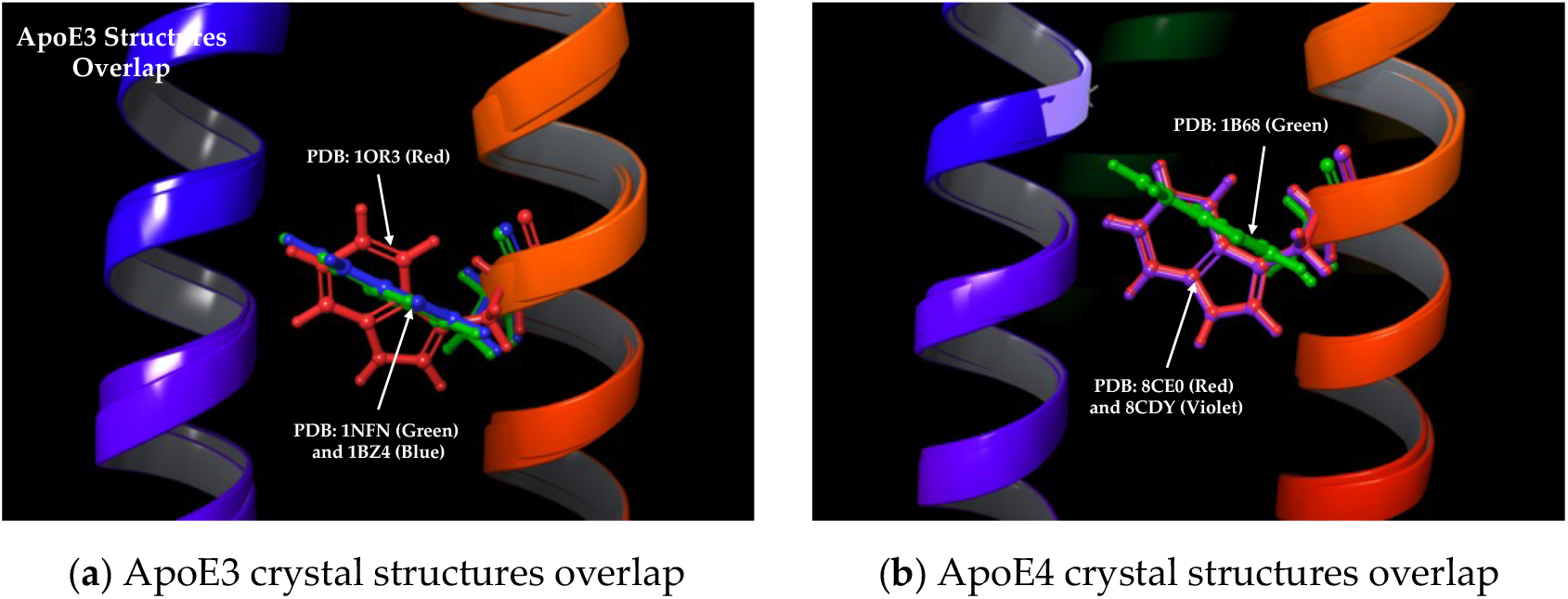
Comparison of ApoE3 and ApoE4 structures: (**a**) Two ApoE3 structures (PDB IDs: 1NFN, 1BZ4) have Tryptophan34 sidechain in expected “flip-in” orientation (green and blue), while one structure (PDB ID: 1OR3) in “flip-out” position (red) resembling ApoE4; (**b**) Two ApoE4 structures (PDB IDs: 8CE0, 8CDY) with Tryptophan34 sidechain in expected “flip-out” position (red and violet), while one structure (PDB ID: 1B68) in “flip-in” orientation (green) similar to ApoE3.

### 2.2. Structure-based ensemble virtual screening utilizing complementary consensus scoring

The ensemble molecular docking studies of bioactive natural compounds from the DCABM-TCM database together with the known ApoE4 stabilizers reported by Petros et al. [25] were carried out against the Tryptophan34 binding pockets of all six ApoE3/E4 structures. The AutoDock Vina [30] docking algorithm was used, whereby protein remains rigid while allowing the ligand flexibility. The AutoDock Vina correctly predicted the binding modes of two available crystal ligands (PubChem CIDs: 83673143 and 137796780) with respective ApoE4 structures (PDB IDs: 6NCN and 6NCO). As seen, there was a good overlap between the ligand binding mode observed by X-ray crystallography and the binding mode predicted by AutoDock Vina (**Figure 3**).

**Figure 3.**
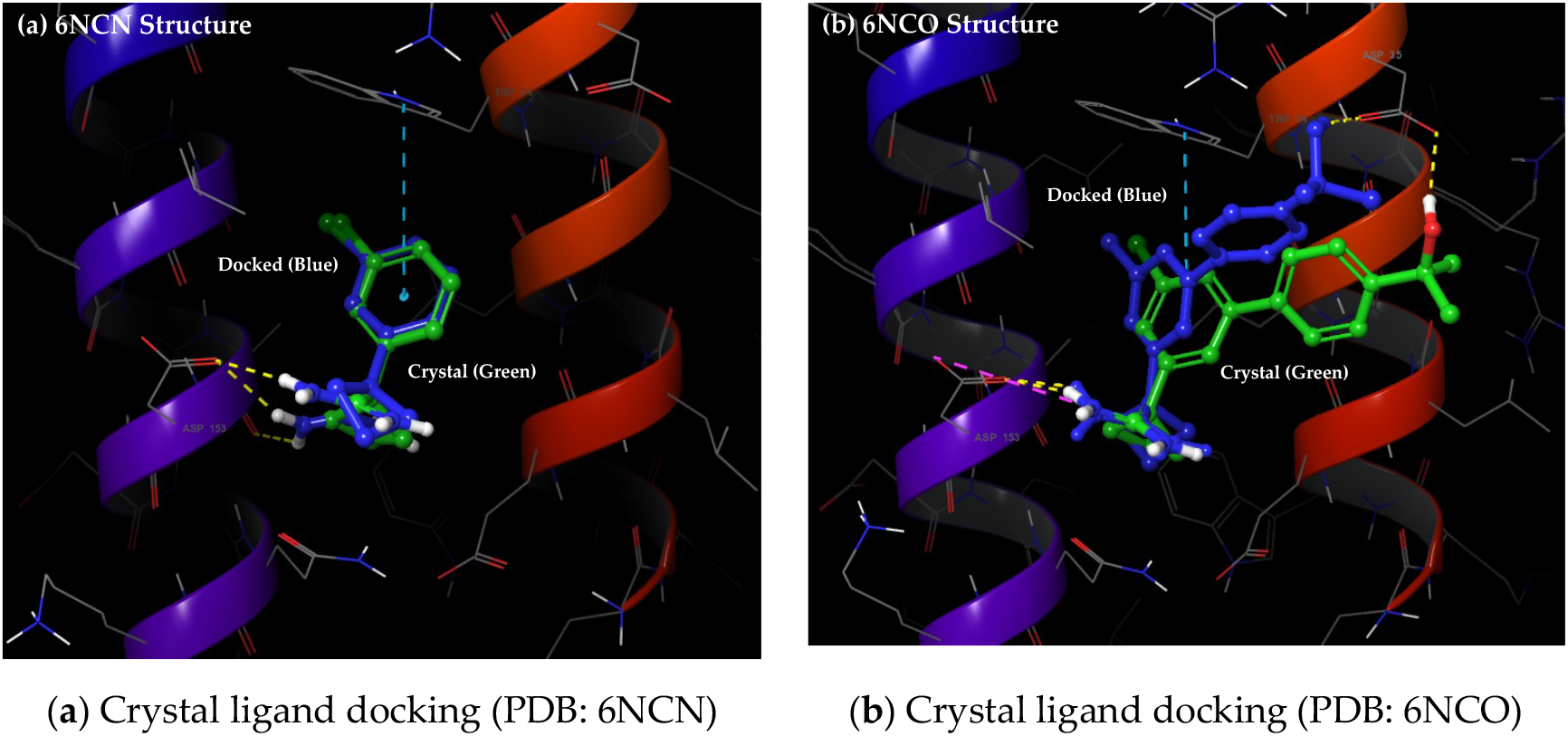
Crystal ligand re-docking: (**a**) PDB ID: 6NCN (ligand PubChem CID: 83673143); (b) PDB ID: 6NCO (ligand PubChem CID: 137796780). Crystal ligands (green carbons) and docked (blue).

Importantly, the docked ligands replicated important ligand-protein interactions, including pi-pi interaction with Tryptophan34 and H-bond interactions with Aspartate35 and 153 residues. Thus, AutoDock Vina proved well-suited for our ensemble docking studies against ApoE proteins and revealed varying binding ranks for the known ApoE4 ligands (**Table 1**).

**Table 1.** The AutoDock Vina binding ranks for known ApoE4 ligands against ApoE3, free-ApoE4 and holo-ApoE4.

| PubChem CID | NMR Kd (μM) | ApoE3 (1NFN) | ApoE3 (1BZ4) | ApoE4 (8CE0) | ApoE4 (8CDY) | Holo-ApoE4 (6NCN) | Holo-ApoE4 (6NCO) | Consensus Rank |
| --- | --- | --- | --- | --- | --- | --- | --- | --- |
| 83673143 | 900 ± 108 | 1027 | 1013 | 1029 | 993 | 733 | 853 | 286 |
| 155530661 | 230 ± 24 | 328 | 490 | 185 | 204 | 519 | 572 | 1167 |
| 137796780 | 30 ± 4 | 527 | 565 | 242 | 298 | 65 | 45 | 28 |
| 155563897 | 20 ± 3 | 426 | 208 | 186 | 628 | 60 | 18 | 9 |
| 155552638 | 20 ± 4 | 444 | 400 | 660 | 664 | 109 | 76 | 14 |
| 155557185 | 8 ± 3 | 471 | 652 | 519 | 505 | 61 | 25 | 6 |
| 155511476 | < 5 | 470 | 651 | 827 | 774 | 111 | 95 | 11 |
| 155538646 | < 5 | 594 | 611 | 659 | 744 | 116 | 148 | 21 |
| Correlation Score <sup>1</sup> |  | 0.839 | 0.723 | -0.546 | -0.738 | 0.953 | 0.973 | **0.996** |

<sup>1</sup> Correlations scores between AutoDock Vina ranks and experimental Kd values for ApoE4 binding.

Several key observations were made based on these ensemble docking data against ApoE3 and E4 structures. Specifically, all these know ApoE ligands, except the smallest fragment (PubChem CID: 83673143, MW: 208.69 g/mol), showed varying degrees of binding affinity across ApoE3, free-ApoE4 and holo-ApoE4 structures. The ApoE3 and holo-ApoE4 structures, with similar Tryptophan34 sidechain orientation (**Figure 1**), present similar binding pockets, but only the holo-ApoE4 showed excellent correlation between the AutoDock Vina ranks and the experimental NMR Kd values reported by the AbbVie team [25]. Furthermore, free-ApoE4 structures (8CE0 and 8CDY) exhibited negative correlation scores as opposed to positive correlation scores observed with holo-ApoE4 structures (6NCN and 6NCO). This suggested that molecules that bind well with holo-ApoE4 pocket bind poorly with free-ApoE4 structure. This is in agreement with the “flip-in” Tryptophan34 orientation in holo-ApoE4 structures (**Figure 1**) that presents a more open, well-defined binding pocket, as opposed to “flip-out” Tryptophan34 orientation in free-ApoE4 structures closing this pocket. Based on this observation, we devised a new consensus ranking scheme (**Equation 1**) involving AutoDock Vina binding ranks against both the free- and holo-ApoE4 structures.

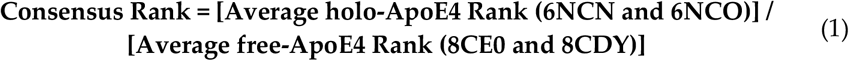

Our novel consensus ranking scheme complemented the docking analysis well and further improved the correlation score in comparison with the experimental Kd values to a near-perfect 0.994. Notably, it also significantly improved the overall docking ranks of the more active ApoE4 ligands (Kd values < 30 µM), putting them in the top ∼1.6% hitlist. Accordingly, the top 25 novel hits out of ∼1,250 DCABM-TCM database molecules (top ∼2%) obtained through this ensemble docking followed by consensus ranking are listed below (**Table 2**).

**Table 2.** Top 25 hits each from the structure-based ensemble docking and shape-based screening.

| Sr. No. | Consensus Docking Hits<br>(PubChem CIDs) | Consensus ROCS Hits<br>(PubChem CIDs) |
| --- | --- | --- |
| 1 | 100781 | 53266 |
| 2 | 161294 | 160487 |
| 3 | 91510 | 5320382 |
| 4 | **193679**<sup>1</sup> | 114829 |
| 5 | 441805 | 5282073 |
| 6 | 11556558 | 632135 |
| 7 | 94320 | **193679**<sup>1</sup> |
| 8 | 9974201 | 46886723 |
| 9 | 10114 | 115067 |
| 10 | 71773126 | 5281601 |
| 11 | 163184367 | 5316844 |
| 12 | 14077830 | 92775 |
| 13 | 9983614 | 5280443 |
| 14 | 11003773 | 96539 |
| 15 | 64945 | 5281617 |
| 16 | 102501232 | 5322078 |
| 17 | 73347309 | 10450045 |
| 18 | 9548703 | 198910 |
| 19 | 14463159 | 5319081 |
| 20 | 91471 | 5320053 |
| 21 | 5380976 | 11601633 |
| 22 | 114850 | 439246 |
| 23 | 163005195 | 5318980 |
| 24 | 92097 | 5281698 |
| 25 | 151529 | 72303 |

<sup>1</sup> The combined docking- and shape-based screening revealed 193679 as a common hit (shown bold).

### 2.3. Ligand-based virtual screening using 3D shapes of most active ApoE4 stabilizers

The ligand-centric, shape-based virtual screening has been suggested to obtain more consistent performance across targets in terms of hit rate than other 3D screening methods including structure-based docking and 3D pharmacophore screening [31–33]. More importantly, shape similarity techniques often lead to new inhibitors with innovative chemical scaffolds. Therefore, we decided to screen the DCABM-TCM database using the **R**apid **O**verlay of **C**hemical **S**tructures (ROCS) method [34] that aligns molecules based on shape and/or chemical similarity (ROCS 3.7.0.0. OpenEye, Cadence Molecular Sciences, Santa Fe, NM. http://www.eyesopen.com). In this regard, two crystal structures of small-molecule stabilizers in complex with ApoE4 are currently available (PDB IDs: 6NCN and 6NCO), which present the tentative spatial and chemical requirements necessary for the effective binding of a given molecule to the Tryptophan34 pocket on ApoE4.

Unfortunately, the two ligands associated with these two crystal structures (PubChem CIDs: 83673143 and 137796780) are relatively weak ApoE4 ligands possessing poor Kd values to ApoE4 (**Table 1**). Therefore, we decided to use the two most active ApoE4 ligands (PubChem CIDs: 155511476 and 155538646) with ApoE4 Kd < 5 µM for our ligand-based screening. The best docking conformations of these two ligands, which recapitulated the favorable H-bond and pi-pi interactions in the Tryptophan34 pocket of ApoE4 protein (PDB ID: 6NCO), were used as the queries for the 3D shape-based ROCS screening (**Figure 4a**). The multi-conformer database of TCM-derived molecules was screened to rank the database molecules using ROCS_TanimotoCombo (shape and color) score of the highest-ranking conformer of each compound. These shape-similarity scoring data obtained using two known ApoE4 ligands were then combined to reveal top 25 molecules with the best ROCS_TanimotoCombo scores (**Table 2**). As seen, a natural flavonoid Isobavachin (PubChem CID: 193679) was revealed as the common hit from both the structure- and ligand-based screenings, possessing high docking and 3D shape ranks. The 3D molecular structure and shape of Isobavachin (**Figure 4b**) mimic those that of two query ApoE4 ligands (**Figure 4a**), with ROCS_TanimotoCombo score of 0.904 and possessing key H-bond capabilities on two ends. Specifically, the amine groups in the known ApoE4 ligands are mimicked by the hydroxyl groups in Isobavachin. Furthermore, a hydrophobic Cl in known ApoE4 ligands is observed to occupy the Tryptophan34 pocket in ApoE4, which is mimicked by the hydrophobic prenyl group in Isobavachin. Thus, Isobavachin appears to be a top promising hit molecule, presenting a novel scaffold as compared to the known ApoE4 ligands (Tanimoto Similarity 0.03 and 0.04 compared to 155511476 and 155538646, respectively).

**Figure 4.**
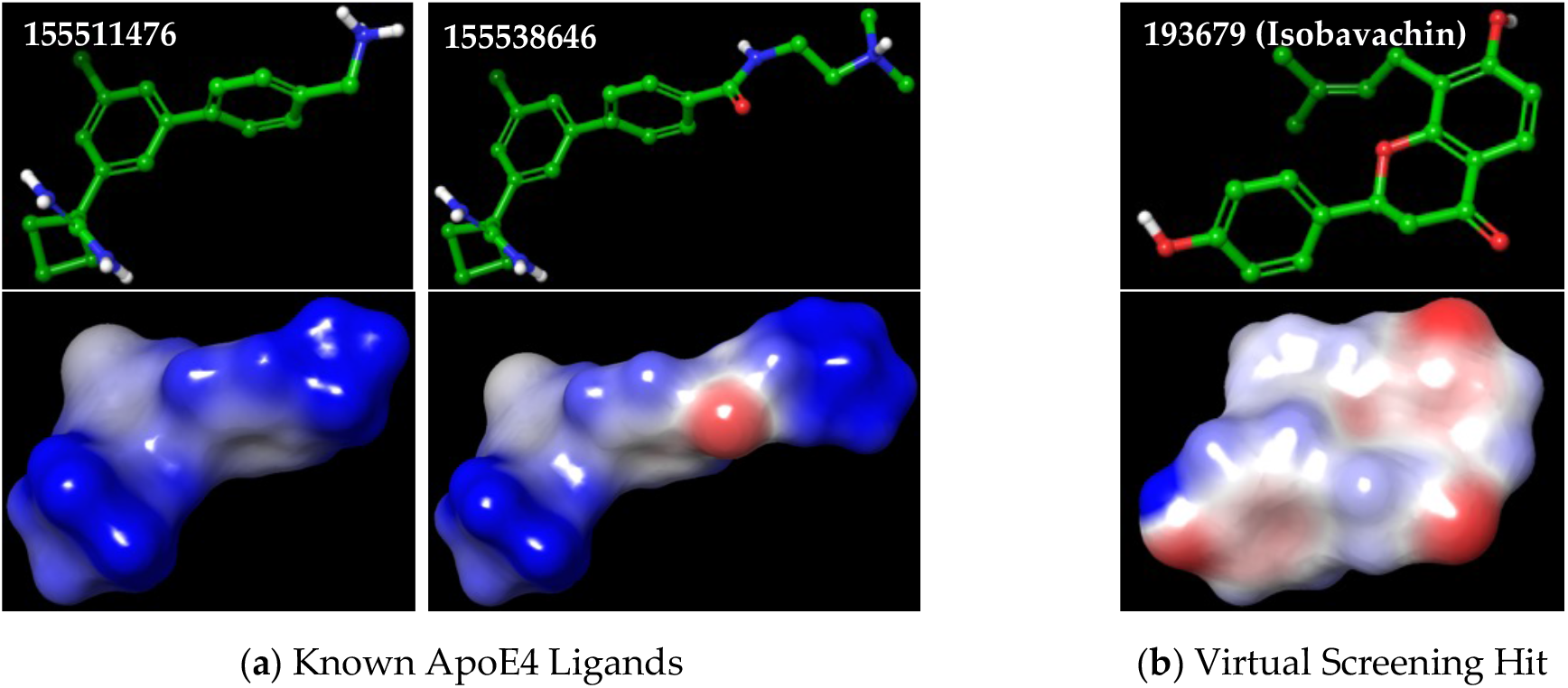
The 3D molecular structures and shapes of known ApoE4 ligands and our top hit molecule: (**a**) Two most active ApoE4 ligands (PubChem CIDs: 155511476 and 155538646); (**b**) Top virtual screening hit Isobavachin (PubChem CID: 193679).

### 2.4. Flexible receptor docking to investigate ligand-induced flipping of Tryptophan34 residue

The molecular modeling studies of total 49 hits from docking and shape screening together with 8 known ApoE4 stabilizers from AbbVie [25] were carried out for investigating their binding modes within the Tryptophan34 pocket on free-ApoE4 (PDB ID: 8CE0) that has Tryptophan34 residue in “flip-out” orientation (**Figure 1**). All these hit compounds are structurally different and thus may not be expected to fit in the ApoE4 pocket in the same way as that of the known ligands. Therefore, the **I**nduced-**F**it **D**ocking (**IFD**) methodology was used for docking, as it has been shown to be efficient in handling the receptor flexibility for structurally different ligands [35,36]. The focus was on the ligands capable of replicating the ligand-protein interactions exhibited by the known ApoE4 stabilizers. Specifically, H-bond interactions with Aspratate35 and 153, and most importantly flipping of the Tryptophan34 residue to “flip-in” position, thus opening up a pocket that is to be filled by a suitable hydrophobic group. The IFD studies were successful with all the 8 known ApoE4 ligands successfully changing the orientation of the Tryptophan34 sidechain from the “flip-out” position in free-ApoE4 to the “flip-in” position after ligand binding (**Table 3**). Notably, even the smallest fragment (CID: 83673143) with molecular weight of 208.69 g/mol successfully flipped the Tryptophan34 orientation to resemble that of ApoE3, thus further emphasizing the suitability of the IFD protocol used here. The other desired ligand-protein interactions varied across different ligands, correlating well with their reported experimental binding Kd values. For example, top 6 out of 8 compounds with Kd values ≤ 30 µM exhibited all the key interactions, while the two weakest molecules lacked key H-bond interactions with either Aspartate35 or 153, further highlighting their importance.

**Table 3.** The ligand-protein interaction characteristics obtained for the IFD binding modes of known ApoE4 ligands.

| PubChem CID | NMR Kd (μM) | Tryptophan34 orientation | Aspartate35 Interaction | Aspartate153 Interaction | Hydrophobic Pocket Filling |
| --- | --- | --- | --- | --- | --- |
| 83673143 | 900 ±108 | Flip-in | — | H-bond | Cl atom |
| 155530661 | 230 ± 24 | Flip-in | H-bond | — | Cl atom |
| 137796780 | 30 ± 4 | Flip-in | H-bond | H-bond | Cl atom |
| 155563897 | 20 ± 3 | Flip-in | H-bond | H-bond | Cl atom |
| 155552638 | 20 ± 4 | Flip-in | H-bond | H-bond | Cl atom |
| 155557185 | 8 ± 3 | Flip-in | H-bond | H-bond | Cl atom |
| 155511476 | < 5 | Flip-in | H-bond | H-bond | Cl atom |
| 155538646 | < 5 | Flip-in | H-bond | H-bond | Cl atom |

In this context, out of 49 virtual hits from docking and shape-based screening, only the top common hit Isobavachin exhibited all the key interactions (**Figure 5**).

**Figure 5.**
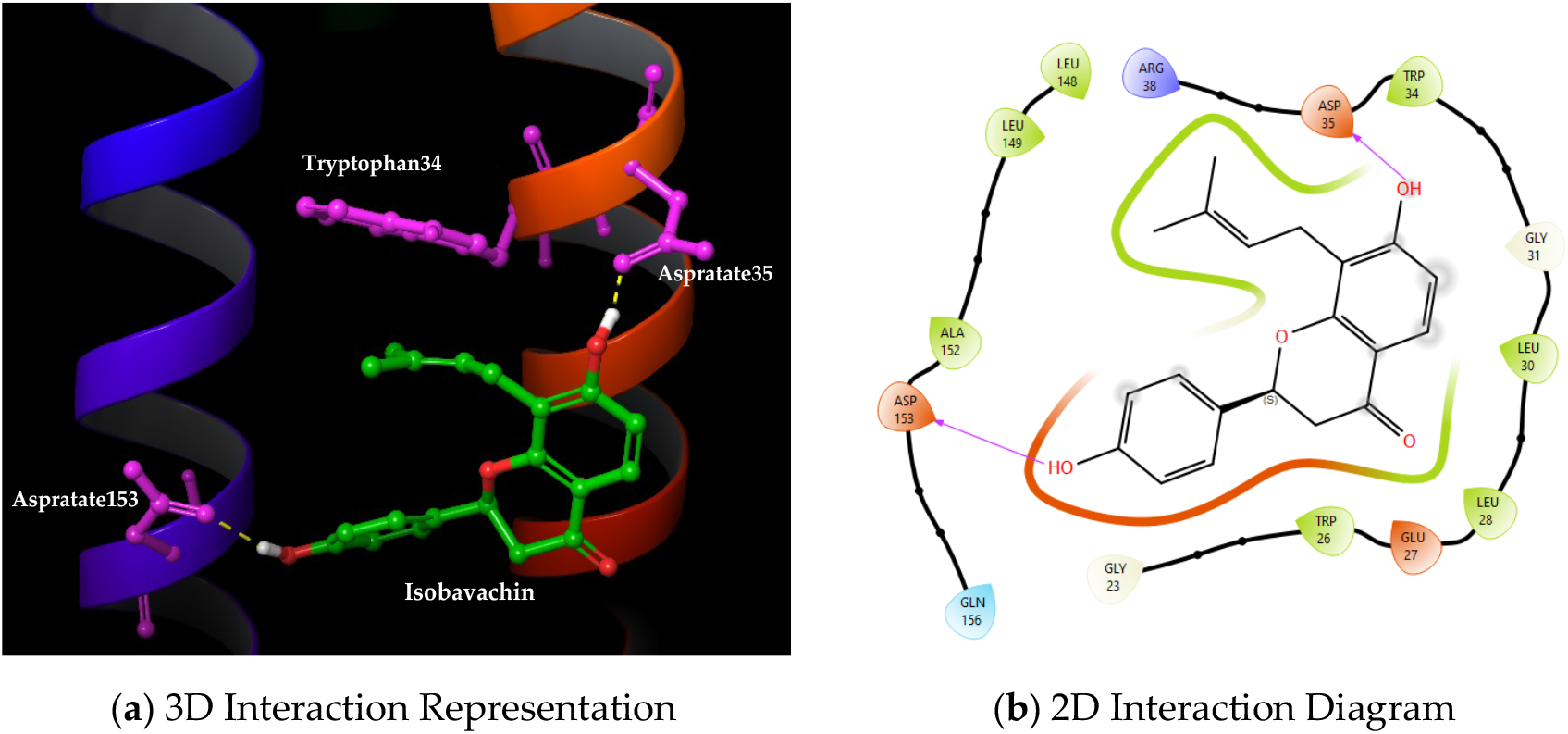
The binding mode of the top-ranking hit molecule Isobavachin (PubChem CID: 193679) obtained through the induced-fit docking: (**a**) Tryptophan34 binding pocket on ApoE4 containing bound ligand; (**b**) 2D ligand–protein interaction diagram.

Notably, Isobavachin flipped the Tryptophan34 sidechain to the desired “flip-in” position, formed critical H-bond interactions with Aspartate35 and 153 residues, and also its hydrophobic prenyl perfectly occupied the Tryptophan34 binding pocket on ApoE4. This IFD binding mode of Isobavachin is in accordance with those of the known ApoE4 stabilizers from AbbVie, especially the most active ones with Kd values ranging from ∼ 5 – 30 µM. Thus, Isobavachin possesses key structural features required to occupy the Tryptophan34 binding pocket on ApoE4, potentially leading to its structure correction and stabilization resembling to the ApoE3 protein.

### 2.5. Molecular dynamics simulation of Isobavachin-ApoE4 complex

The stability of our top virtual hit Isobavachin within the ApoE4 binding pocket was further investigated by carrying out molecular dynamics (MD) simulations. The root-mean-square-distance (RMSD) analysis of Isobavachin over 100 ns MD simulation revealed its stable binding in the Tryptophan34 pocket on ApoE4 (**Figure 6a**). After initial configuration change, Isobavachin showed consistent RMSD value around ∼ 2.0 Å throughout the 100 ns simulation period, thus indicating its stable binding with the ApoE4 pocket. Furthermore, RMSD profile for Isobavachin-bound ApoE4 backbone was < 2.7 Å throughout the simulation, thus further supporting stability of the ligand-protein complex. Indeed, Isobavachin showed relatively much better binding stability with the ApoE4 pocket as compared to the two most active ApoE4 stabilizers reported by AbbVie (CIDs: 155511476 and 155538646). Accordingly, Isobavachin showed consistently favorable interaction profile with key amino acids such as Tryptophan34, Aspartate35 and 153, throughout the 100ns simulation interval (**Figure 6b**).

**Figure 6.**
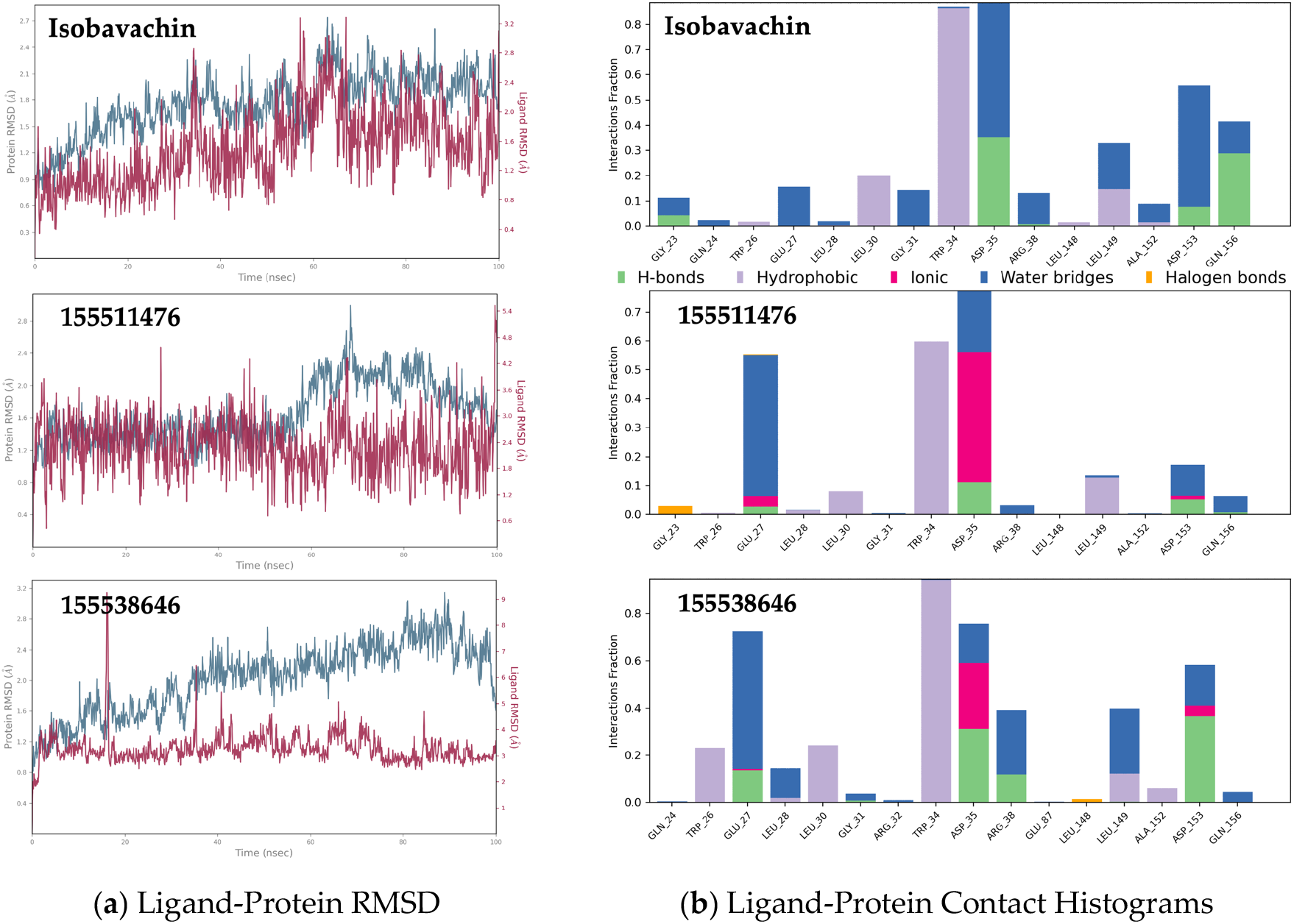
The MD simulation data: (a) The RMSD analysis over 100ns MD simulation of ligand binding with the Tryptophan34 pocket on ApoE4; (b) The ligand-protein interaction histogram.

Notably, Isobavachin effectively changed the orientation of the Tryptophan34 sidechain in ApoE4 to the desired “flip-in” position (**Figure 7a**) and most importantly, successfully maintained this orientation throughout the 100 ns MD simulation (**Figure 7b**). As seen, the orientation of Tryptophan34 sidechain residue averaged over the simulation trajectory remained parallel to the protein axis in free-ApoE4, while flipped perpendicular to the protein axis in Isobavachin-bound ApoE4, making it similar to ApoE3. Thus, Isobavachin seems to be an ideal drug candidate for correcting the pathological ApoE4 structure to the beneficial ApoE3 structure.

**Figure 7.**
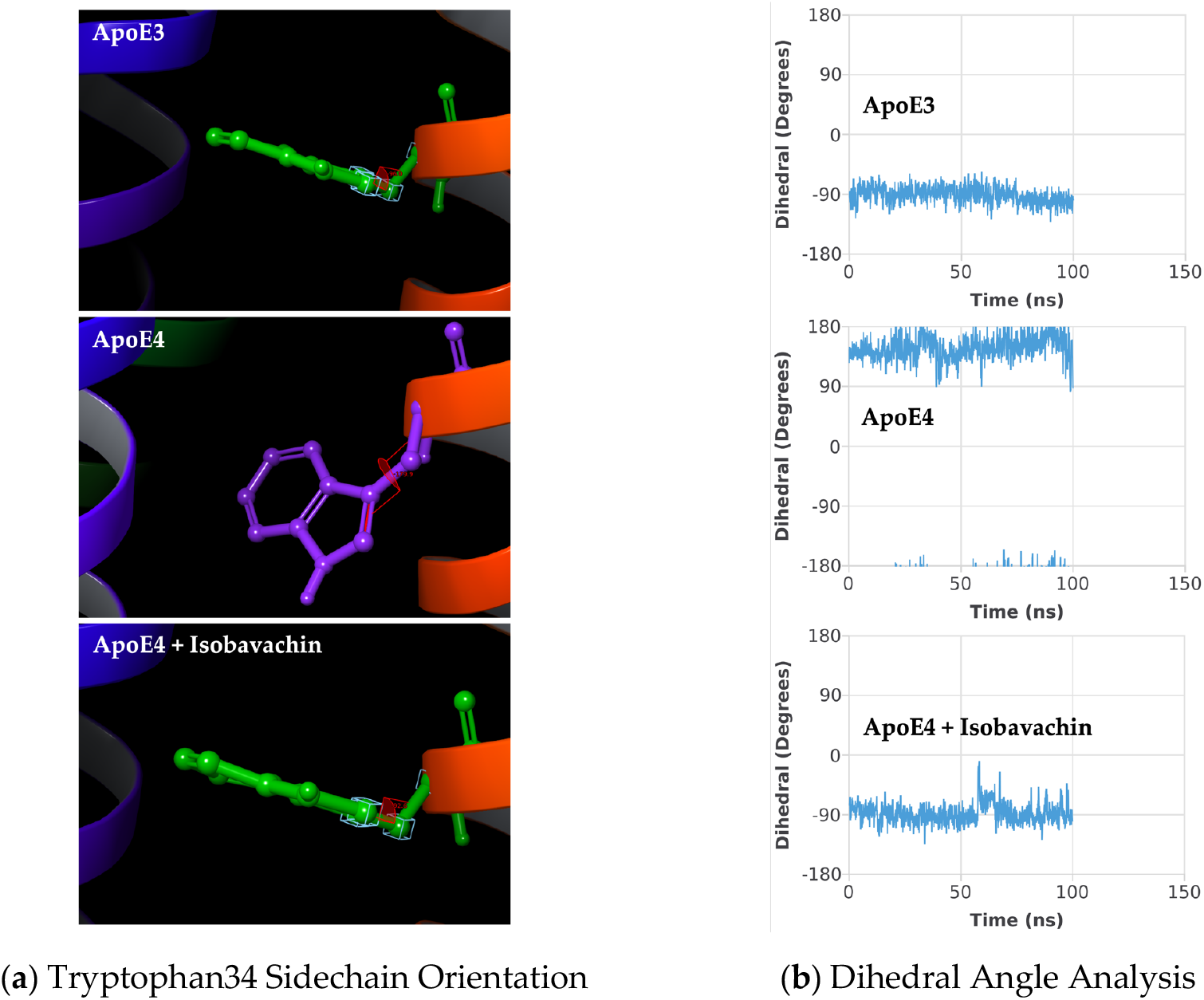
The Tryptophan34 sidechain flipping: (a) The ApoE protein structures averaged across 100 ns MD simulation, showing the representative Tryptophan34 sidechain orientation; (b) The dihedral angle analysis for the Tryptophan34 residue.

### 2.6. Biophysical binding analysis of Isobavachin and related flavonoids with ApoE3 and ApoE4

The top virtual hit Isobavachin was further subjected to the Surface Plasmon Resonance (SPR) analysis using Biacore T200 to determine its binding constants with both the ApoE3 and ApoE4 proteins. Together with the human recombinant ApoE3 and ApoE4 proteins, a negative control protein 2’-deoxynucleoside 5’-phosphate N-hydrolase (His-Sumo-DNPH1) was also included in the SPR analysis. The His-Sumo-DNPH1 is structurally unrelated to ApoE proteins, while having similar size as ApoE proteins (∼ 30 KDa). In agreement with our extensive computational analysis, Isobavachin showed strong binding with the ApoE4 protein with the Kd value of ∼ 540 nM (**Figure 8**).

**Figure 8.**
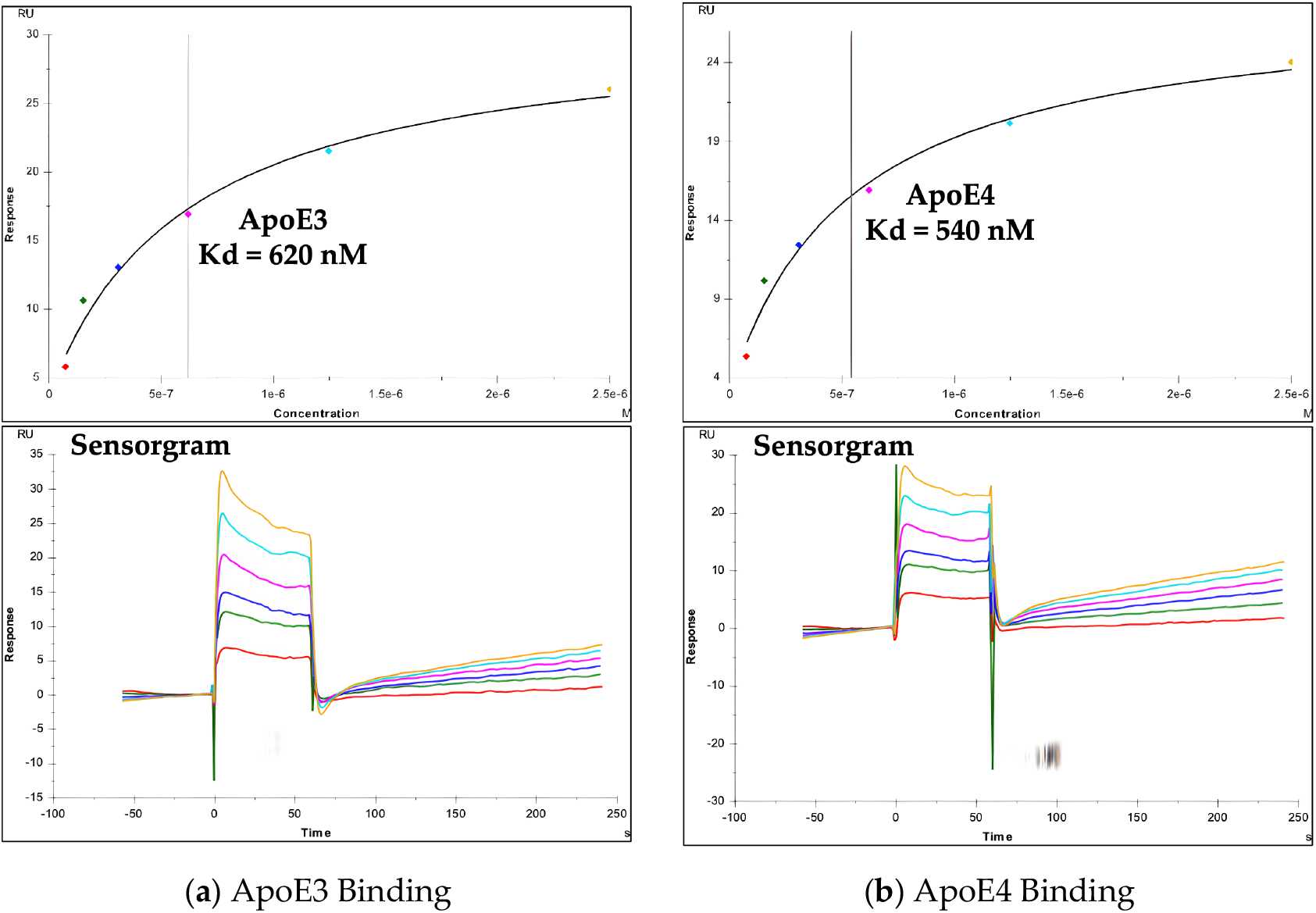
The Surface Plasmon Resonance (SPR) analysis: (a) Isobavachin binding with human recombinant ApoE3; (b) Isobavachin binding with human recombinant ApoE4.

Isobavachin also showed similar Kd value with ApoE3 (∼ 620 nM), thus further supporting potential re-orientation of Tryptophan34 sidechain residue in ApoE4 resembling that in ApoE3, presenting similar binding pockets for Isobavachin. In contrast, Isobavachin did not bind well with the negative control protein His-Sumo-DNPH1 (Kd value > 100 µM), thus emphasizing selective binding to the ApoE proteins (**Table 4**).

**Table 4.**
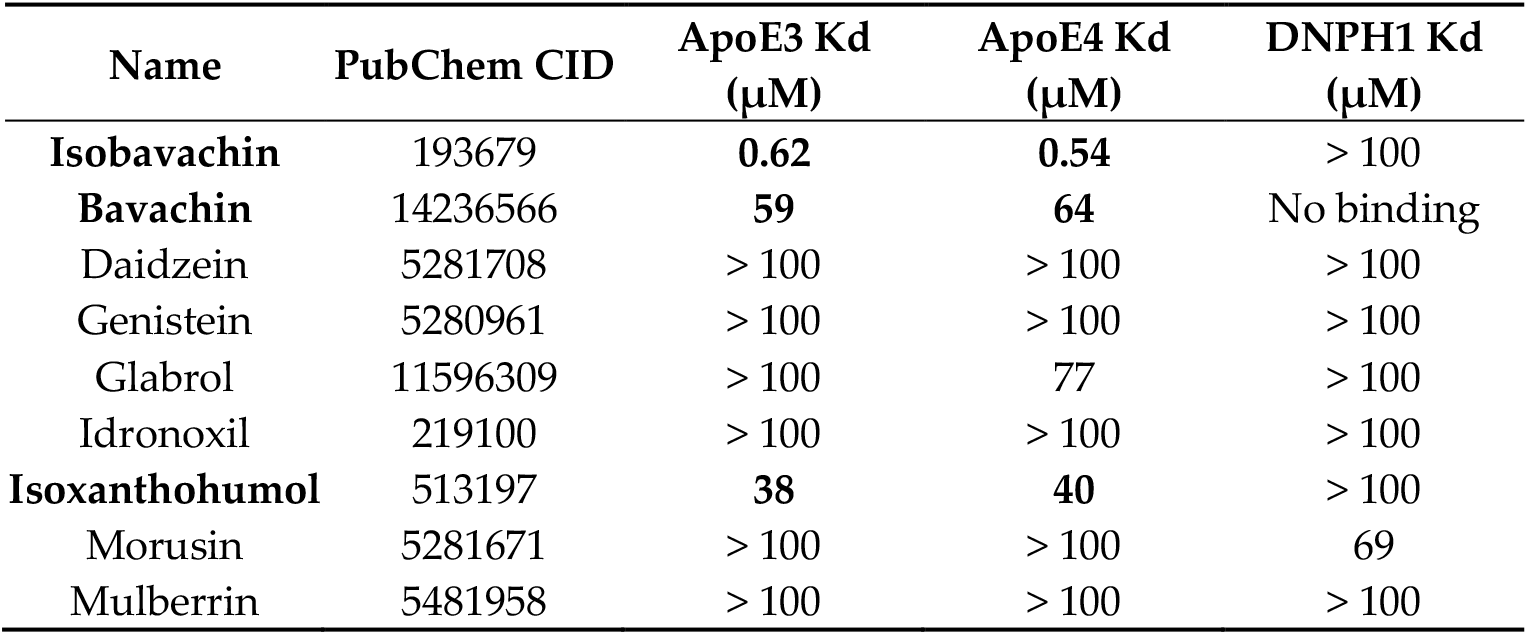
The SPR binding data for Isobavachin and other bioactive flavonoids.

In addition to Isobavachin, we also tested a few other bioactive, purchasable flavonoids for their potential binding with the ApoE3/4 proteins. These included various prenylated (Bavachin, Glabrol, Isoxanthohumol, Morusin, Mulberrin) and non-prenylated (Daidzein, Genistein, Idronoxil) flavonoids (**Figure 9**).

**Figure 9.**
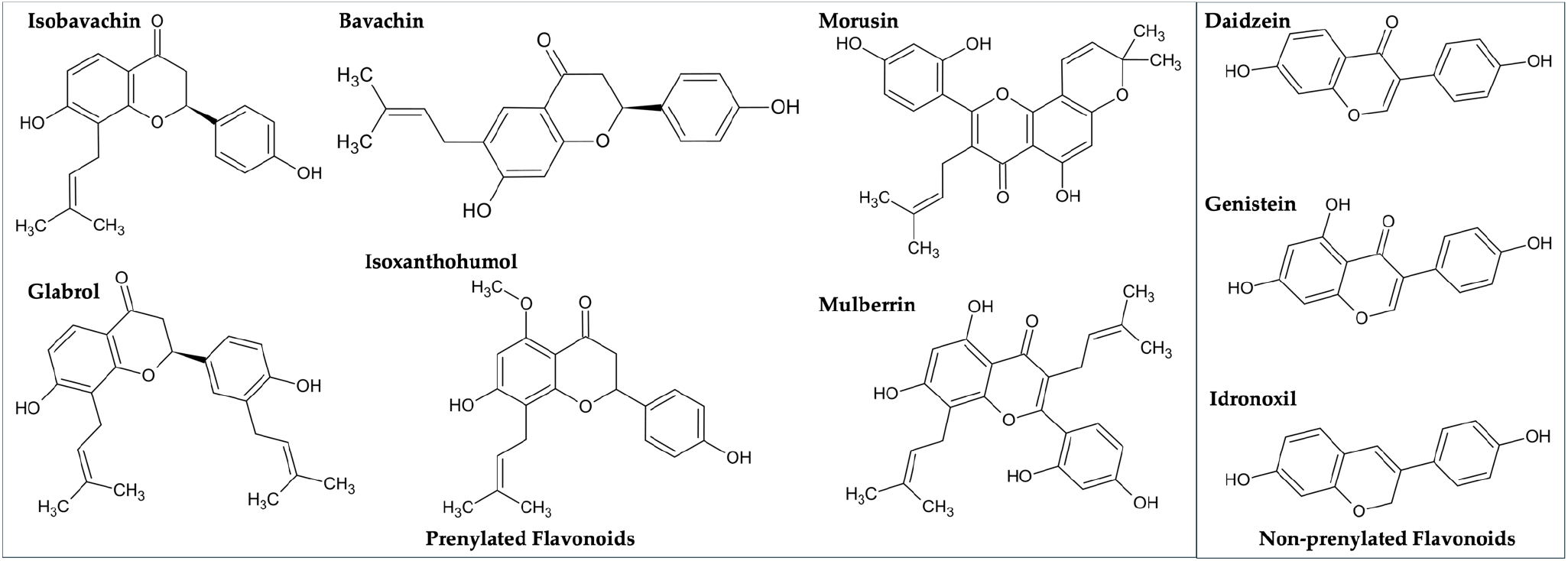
The chemical structures of Isobavachin and other flavonoids tested in the present study.

Only two of these flavonoids, viz. Bavachin and Isoxanthohumol, both prenylated like Isobavachin, showed some binding activity against ApoE3 and ApoE4 proteins (**Table 4**). Overall, our top virtual hit Isobavachin proved to be the strongest ApoE3/E4 binder.

## 3. Discussion

The present study, through a robust combination of structure- and ligand-based virtual screening methodologies, effectively highlights the potential of the natural medicinal compounds from the DCABM-TCM as a rich repository of bioactive compounds for AD drug discovery. The identification of Isobavachin as a natural flavonoid capable of correcting and potentially stabilizing the ApoE4 structure marks a significant step forward in AD research.

ApoE plays a central role in AD pathogenesis, with the “ApoE Cascade Hypothesis” postulating that the isoforms of ApoE (E2, E3, and E4) exert differential effects on AD pathology through complex molecular and cellular mechanisms, contributing to the varying levels of risk and progression of the disease [37]. ApoE4, the strongest genetic risk factor for late-onset AD, promotes Aβ aggregation and deposition, disrupts lipid homeostasis, and exacerbates neuroinflammation and synaptic dysfunction. Mechanistically, ApoE4 has been shown to be less efficient in clearing Aβ compared to ApoE2 and ApoE3, leading to its accumulation in the brain [38]. Furthermore, ApoE4 undergoes proteolytic cleavage, generating toxic fragments that impair mitochondrial function and neuronal viability [39]. This cascade of events accelerates tau pathology and neurodegeneration, creating a feedback loop that perpetuates AD progression [40]. Therapeutically, targeting the ApoE cascade, through stabilizing ApoE4 structure (both domain interaction and Trptophan34 sidechain orientation), modulating its lipidation, or mitigating its pathological interactions, offers promising avenues for curative interventions in AD [19,25,41,42].

By stabilizing the ApoE4 structure through Tryptophan34 re-orientation, Isobavachin may mitigate its toxic gain-of-function properties and effectively reduce its contribution to neurodegenerative processes in AD. Importantly, Isobavachin is a bioactive compound, a naturally occurring prenylated flavonoid derived primarily from the seeds of Psoralea corylifolia (commonly known as Babchi). Prenylated flavonoids have been shown to exhibit wide-ranging medicinal properties, such as anti-oxidant, anti-inflammatory, anti-cancer, anti-diabetic, anti-microbial, and neuroprotective effects, making it an attractive candidate for therapeutic applications [43]. It is noteworthy that Isobavachin has been shown to readily penetrate the blood-brain barrier (BBB), thereby enabling its potential to directly affect central nervous system (CNS) pathologies including AD [44]. Overall, the natural origin of Isobavachin and its bioavailability as a TCM-derived compound, coupled with its ApoE4-centric effects outlined in this study, underscores its translational potential as a favorable therapeutic agent against AD.

Our combined molecular docking and 3D shape screening protocol successfully identified Isobavachin as the top hit which was then experimentally confirmed to be a potent ApoE ligand. Importantly, the present study underscores the importance of selecting appropriate protein structures for virtual screening of potential ApoE4 stabilizers that act through ligand-induced conformation changes. Our decision to exclude atypical ApoE structures based on Trypto-phan34 sidechain orientation resulted in a refined dataset of ApoE3 and ApoE4 structures. This approach in turn ensured the enhanced reliability of computational predictions, as supported by strong correlations between experimental and predicted binding affinities for known ApoE ligands (**Table 1**). Moreover, our novel consensus ranking scheme (**Equation 1**) further improved predictive accuracy by emphasizing the distinct binding properties of free- and holo-ApoE4 structures, yielding a near-perfect correlation score of 0.994 with the experimental Kd values. These encouraging results demonstrate that the adopted integrated virtual and experimental protocol together with our innovative consensus ranking scheme may prove to be a useful tool for identification of novel hits that can be developed into potent ApoE-targeted therapies for AD.

In this context, further experimental validation of Isobavachin’s efficacy in cellular and *in vivo* models of AD is warranted to further confirm its therapeutic potential against various AD-related changes, including Aβ deposition, NFT formation and region-specific neurodegeneration [45,46]. In addition, the medicinal chemistry structural optimizations to enhance binding affinity and specificity for the Tryptophan34 pocket could also be explored to develop novel analogs of Isobavachin. For example, possible replacement of the prenyl group in Isobavachin with other hydrophobic groups may be explored for compounds with improved drug-like properties. Moreover, the replacement of hydroxyl groups with other H-bonding moieties such as amine groups may lead to compounds with improved bioavailability, metabolic stability, receptor binding affinity, and overall therapeutic efficacy [47]. Accordingly, the future *in vivo* studies exploring pharmacokinetics, pharmacodynamics, and safety profiles of Isobavachin and its potent analogs will be crucial for their advancement into preclinical and clinical development.

In summary, the findings of this study establishing Isobavachin as a potential ApoE4 structure corrector may offer significant implications for the development of disease-modifying therapies for AD.

## 4. Conclusions

The present study demonstrates the discovery of Isobavachin, a natural flavonoid, as a promising ApoE4 structure corrector for Alzheimer’s disease (AD). Through an integrated computational and experimental approach, we identified Isobavachin as a top hit that effectively binds to the Tryptophan34 pocket of ApoE4 and induces the desired conformational shift towards an ApoE3-like structure. The observed ligand-induced flipping of the Tryptophan34 sidechain, along with key protein-ligand interactions, highlights the potential of Isobavachin in addressing the aberrant structure and neurotoxic downstream effects of ApoE4.

This work underscores the utility of natural products, particularly flavonoids, as valuable scaffolds for AD drug discovery. By employing complementary structure-based docking, ligand-based virtual screening, and molecular dynamics simulations, followed by biophysical validation, we have established a comprehensive framework for identifying and characterizing ApoE4 stabilizers. Importantly, the study validates the viability of the combined screening protocol to explore other bioactive compound libraries for novel ApoE4-correcting small molecules.

The results lay a strong foundation for further preclinical investigations into Isobavachin and similar flavonoids as potential disease-modifying agents. Future studies will focus on optimizing the chemical scaffold of Isobavachin to enhance its potency, selectivity, and pharmacokinetic properties. Additionally, *in vivo* validation in AD models is necessary to evaluate its therapeutic efficacy and safety profile. By targeting ApoE4’s structural instability, this research opens new avenues for the development of effective and targeted treatments for AD, addressing a critical unmet need in the field.

## 5. Materials and Methods

### Ensemble Molecular Docking

The DCABM-TCM database was prepared for docking by extracting SMILES from: http://bio-net.ncpsb.org.cn/dcabm-tcm/ [26]. The database included ∼1,740 bioavailable TCM compounds detected in the blood, out of which ∼ 500 molecules with high molecular weights were removed. The 3D conformers were generated for the remaining molecules using OMEGA 6.0.0.0 3D conformer generator [48] from OpenEye, Cadence Molecular Sciences, Santa Fe, NM, http://www.eyesopen.com. The ensemble docking of this drug database was carried out using two each crystal structures of the ApoE3 (PDB: 1NFN, 1BZ4)), free-ApoE4 (PDB: 8CE0, 8CDY), and holo-ApoE4 (PDB: 6NCN, 6NCO) proteins downloaded from the protein data bank (PDB) [49]. These ApoE protein structures were processed using the AutoDock Tools utility [50], whereby bound ligands and water molecules were removed, polar hydrogens were added, non-polar hydrogens were merged, and Gasteiger charges were assigned for all the atoms in the proteins. The ApoE proteins and ligand atoms were converted into PDBQT format. The AutoDock Vina docking algorithm [30] was then used to carry out structure-based docking of the DCABM-TCM molecules to the Tryptophan34 binding site on ApoE proteins. All the ApoE structures were overlapped on to the 6NCO crystal structure and thus the same search space coordinates were used for docking into all these ApoE structures [Center-X: 2.30, Y: -12.59, Z: 11.90; Size-X: 25 Å, Y: 25 Å, Z: 25 Å]. Default docking parameters were used, and the docked ligands were ranked according to their best docking score values. A novel consensus scoring utilizing docking scores against both the free- and holo-ApoE4 structures was used to shortlist top 25 hits for further analysis.

### Ligand-Based Virtual Screening

The ligand-based virtual screening was performed using ROCS 3.4.1.0 3D shape-based method [34] from Open-Eye, Cadence Molecular Sciences, Santa Fe, NM, http://www.eyesopen.com. The DCABM-TCM database was processed using OMEGA 6.0.0.0 using default parameters, which generated ∼ 65K 3D conformers. The ROCS was used to explore this 3D multi-conformer database for molecules with similar shape and color as the best docked conformations of the two most active ApoE4 stabilizers reported by AbbVie Pharmaceuticals (PubChem CIDs: 155511476 and 155538646). The database compounds were ranked by the TanimotoCombo scores for their highest-ranking conformers. The 3D shape-similarity scoring data thus obtained across these two known ApoE4 ligands were then combined to short-list top 25 molecules with best TanimotoCombo scores.

### Induced-Fit Docking (IFD)

The flexible-receptor docking of top structure- and ligand-based virtual screening hits was carried out within the Tryptophan34 residue site of the free-ApoE4 protein crystal structure (PDB ID: 8CE0). The IFD protocol was run with default parameters from the Schrodinger’s Maestro graphical user interface. The first stage of IFD protocol involved extended sampling using Glide docking algorithm to generate 80 initial poses through softened-potential docking step, which allows tolerance of more steric clashes as compared to a normal, rigid-receptor docking protocol. The second stage then involved protein refinement using Prime module to allow the active site conformational change around 10 Å of the initial ligand poses. Herein, the energy minimization was carried out using OPLS4 force field and Prime’s implicit solvent model. Finally, the active site was further refined by Prime followed by redocking by Glide XP scoring function leading to the generation of the final IFD models. The top 10 IFD poses for each molecule were graphically visualized using Maestro and the best conformation for each molecule based on the observed binding mode and the IFD score was chosen for further analysis including molecular dynamics (MD) simulations as described below.

### Molecular Dynamics Studies

The binding stability of our top hits was investigated using the molecular dynamics (MD) simulations of the respective ligand-protein complexes using Schrodinger’s Desmond MD module as described previously [51]. Briefly, the top binding conformations from the induced-fit docking were chosen as the starting conformations for the MD simulations. The ligand interactions were modeled with the OPLS4 force field. The ligand-protein docking complexes were solvated using TIP5P water model in an orthorhombic boundary box of 10 Å size in all three directions, followed by their neutralization with appropriate number of Clions. Also, the salt was added at 0.15 M concentration to simulate the physiological salt concentration. Default parameters in Desmond including OPLS4 force field were utilized to equilibrate this molecular system. Finally, each of the equilibrated systems was subjected to 100 ns of MD simulations at constant temperature and pressure of 300 K and 1 atm, respectively. The atom coordinates were recorded at every 100 ps for the follow-up simulation analyses. The root mean square deviation (RMSD) for the proteins and the docked ligands were calculated over the entire simulation trajectory with reference to their respective first frames. The protein interactions with the ligand were normalized over the course of the 100 ns trajectory, with the score value depicted in the stacked bar chart indicating the percentage of the simulation time a specific interaction is maintained. The representative structures averaged over the entire simulation time were obtained and variance of dihedral angles for Tryptophan34 sidechain residue was plotted across the simulation trajectory.

### Surface Plasmon Resonance (SPR)

The biophysical binding analysis of ligands with ApoE3 and ApoE4 proteins was carried out using the SPR technique. The human recombinant ApoE3 or ApoE4 proteins were purchased from PeproTech and reconstituted according to the manufacturer’s instructions. The negative control protein was His-Sumo-DNPH1 (2’-deoxynucleoside 5’-phosphate N-hydrolase 1), produced in-house in E Coli by The Wistar Institute, Philadelphia. The proteins were covalently attached to a carboxymethyl dextran sensor chip (CMD700M, from Xantec bioanalytics) using standard amine coupling in HEPES buffered saline. The chip is first conditioned for 180s at 10 µL/min with 1M NaCl, 0.1 mM Na Borate, pH 9.0 and then activated with 250 mM EDC/100 mM NHS for 600s at 10 µL/min. The ApoE3/4 and DNPH1 proteins diluted in 10 mM sodium acetate (pH 5.5 and 4.0, respectively) were injected and immobilized on respective flow cells. After immobilization, remaining activated sites were blocked with 1 M ethanolamine at 10 µL/min for 180s. Test compounds (from Cayman Chemical and MedChemExpress) were first diluted in pure DMSO, and then further diluted in the running buffer without DMSO (10 mM HEPES, pH 7.4, 150 mM NaCl, 0.005% Tween20) to match the DMSO concentrations of the sample and the running buffer. The association and dissociation times were 60s and 120s, respectively, and the flow rate was 30 µL/min. After each compound injection, the needle was washed in 50% DMSO to reduce compound carry-over. Solvent correction cycles were included to correct for bulk refractive index changes associated with DMSO.

## Acknowledgments

The authors thank John Stoddart (Computer Science, Widener University) for his kind help with the GPU-enabled molecular dynamics simulations. The authors also thank Joel Cassel (The Wistar Institute, Philadelphia, PA) for helping with the surface plasmon resonance (SPR) binding studies.

